# Synthetic OCP Heterodimers are Photoactive and Recapitulate the Fusion of Two Primitive Carotenoproteins in the Evolution of Cyanobacterial Photoprotection

**DOI:** 10.1101/102160

**Authors:** Sigal Lechno-Yossef, Matthew R. Melnicki, Han Bao, Beronda L. Montgomery, Cheryl A. Kerfeld

## Abstract

The Orange Carotenoid Protein (OCP) governs photoprotection in the majority of cyanobacteria. It is structurally and functionally modular, comprised of a C-terminal regulatory domain (CTD), an N-terminal effector domain (NTD) and a ketocarotenoid; the chromophore spans the two domains in the ground state and translocates fully into the NTD upon illumination. Using both the canonical OCP1 and the presumably more primitive OCP2 from *Fremyella diplosiphon*, we show that an NTD-CTD heterodimer forms when the domains are expressed as separate polypeptides. The carotenoid is required for the heterodimeric association, assembling an orange complex which is stable in the dark. Both OCP1 and OCP2 heterodimers are photoactive, undergoing light-driven heterodimer dissociation, but differ in their ability to reassociate in darkness, setting the stage for bioengineering photoprotection in cyanobacteria as well as for developing new photoswitches for biotechnology. Additionally, we reveal that homodimeric CTD can bind carotenoid in the absence of NTD, and name this truncated variant the C-terminal domain-like Carotenoid Protein (CCP). This finding supports the hypothesis that the OCP evolved from an ancient fusion event between genes for two different carotenoid-binding proteins ancestral to the NTD and CTD. We suggest that the CCP and its homologs constitute a new family of carotenoproteins within the NTF2-like superfamily found across all kingdoms of life.

## Introduction

Genome sequencing in conjunction with structural characterization has revealed that a relatively small number of conserved domains comprise the enormous diversity of extant proteins (Baron et al., 1991; Orengo and Thornton, 2005; Lees et al., 2016). Many of the domains within this repertoire are capable of physically interacting with another domain, and frequently become covalently linked by gene fusion events into single polypeptides (Enright et al., 1999; Marcotte et al., 1999). It is through this variability of domain interactions that such a large protein diversity has arisen from a limited repertoire of mix-and-match components. Accordingly, using a concept borrowed from engineering, protein domains can be viewed as modules of cellular machinery (Hartwell et al., 1999; Fraser, 2005). While proteins may evolve through point mutations and/or gene duplication, it is through recombination, including domain fusion events, that novel functions and regulatory mechanisms may evolve rapidly (Enright et al., 1999; Marcotte et al., 1999; Kummerfeld and Teichmann, 2005; Bashton and Chothia, 2007).

Plant photoreceptors are typically modular multi-domain proteins composed of a light-sensing domain and an effector domain, and are represented by phytochromes, cryptochromes, phototropins, and UVR8 photoreceptors (Wang, 2015). Relatively recently it has been shown that the Orange Carotenoid Protein (OCP), which governs photoprotection in cyanobacteria, is also a photoreceptor (Wilson et al., 2006). It contains both sensor and effector domains (Leverenz et al., 2014). The modular OCP domain architecture is comprised of a regulatory C-terminal domain (CTD), an effector N-terminal domain (NTD), and a non-covalently bound ketocarotenoid chromophore which spans the two domains. Additionally, a flexible and unstructured linker covalently tethers the two domains as a single polypeptide (Fig. 1)(Kerfeld et al., 2003; Leverenz et al., 2014). The NTD fold is comprised of two discontinuous four-helix bundles and appears to be unique to cyanobacteria (Kerfeld et al., 2003). The fold of the CTD has a superficial resemblance to other photosensory domains, namely LOV and BLUF (Kirilovsky and Kerfeld, 2013), but is classified as a member of the diverse NTF2-like superfamily (PFAM 02136) which includes a variety of enzyme domains that bind oxygenated hydrophobic substrates and is widespread in both Eukaryotes and Prokaryotes.

**Figure 1.**
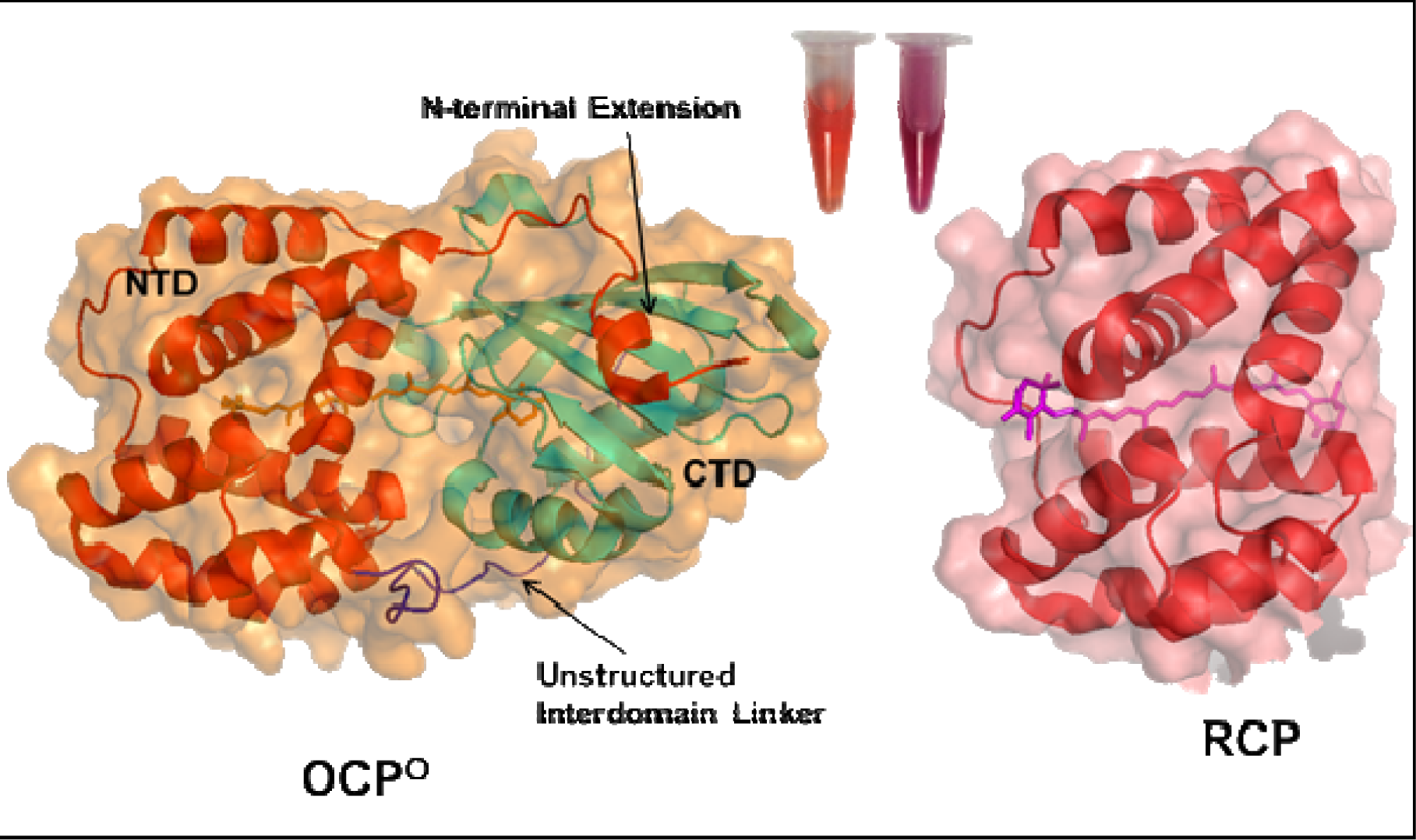
Domain architecture of the Orange Carotenoid Protein. Crystal structures of the *Synechocystis* sp. PCC 6803 OCP^O^ (pdb:4XB5) and its truncated NTD, the Red Carotenoid Protein (RCP) (pdb:4XB4) (Leverenz et al., 2015), illustrating the modular domain architecture and carotenoid translocation. The color of the purified proteins in solution is shown.

In the dark, the NTD and CTD are closely associated (Fig. 1), the protein appears orange (OCP^O^), and it is inactive for phycobilisome (PBS) quenching (Wilson et al., 2006; Wilson et al., 2010). An N-terminal extension from the NTD (NTE) is in direct contact with the CTD (Fig. 1) (Kerfeld et al., 2003). Upon illumination with blue-green light, the hydrogen bonds between the carotenoid and the CTD break, the carotenoid translocates completely into the NTD (Leverenz et al., 2015) and the two domains fully dissociate (Gupta et al., 2015). In this conformation, the protein appears red (OCP^R^) and the NTD is available to interact with the PBS core. The interaction between OCP^R^ and the PBS places the carotenoid in close proximity to bilins in the PBS core, allowing excitation energy (measured as fluorescence) to be quenched and converted into thermal energy (Leverenz et al., 2014; Gupta et al., 2015; Leverenz et al., 2015; Harris et al., 2016). The completion of the OCP photocycle involves a slow dark reversion from OCP^R^ back to OCP^O^ that is accelerated by interaction of its CTD with a second protein, the Fluorescence Recovery Protein (FRP) (Boulay et al., 2010; Gwizdala et al., 2013; Sutter et al., 2013). As such, the OCP is structurally and functionally modular like many photoreceptor proteins; however, its chromophore is needed for both sensory and effector domain functions. Furthermore, in contrast to other photoreceptor proteins, its light-driven domain dissociation is unique.

The first structural characterization of the OCP (Kerfeld et al., 2003) coincided with the beginning of the genomics era, when newly available cyanobacterial genomes revealed that single-domain proteins homologous to either the NTD or to the CTD are encoded in in diverse cyanobacterial genomes (Kerfeld et al., 2003; Kerfeld, 2004a, b; Kirilovsky and Kerfeld, 2012, 2013). The NTD homologs were recently analyzed phylogenetically, classified into several homologous sub-clades, shown to bind carotenoid, and were given the name “Helical Carotenoid Proteins” (HCPs) (Melnicki et al., 2016). Four HCPs have been functionally characterized to date (López-Igual et al., 2016), suggesting the HCP variants are likely to have evolved different roles, with only HCP4 capable of binding and quenching PBS. Homologs to the CTD are also present in cyanobacterial genomes (Kirilovsky and Kerfeld, 2012), and are now termed C-terminal Domain Homologs (CTDHs) (López-Igual et al., 2016; Melnicki et al., 2016). The CTDH genes are very often encoded at loci adjacent to HCP4 or HCP5 genes (Melnicki et al., 2016), leading to a hypothesis that CTDHs might interact with HCPs, and that a domain fusion between CTDH and HCP led to the evolution of the OCP.

A recent phylogenetic analysis using all available cyanobacterial genomes revealed that there are heretofore unknown multiple families of the OCP (Bao et al., submitted). To distinguish these families, the clade containing the well-characterized OCP from *Synechocystis* sp. PCC 6803 was named OCP1. The first characterization of an OCP2, from *Fremyella diplosiphon* UTEX481, demonstrated that OCP2 photoactivation kinetics, transcriptional regulation, oligomeric state, and ability to interact with FRP are distinctly different from its OCP1 paralog (Bao et al., submitted). *Fremyella* OCP2 is photoactivated significantly faster, and reverts back to the orange form almost immediately in the dark, without interaction with FRP. It is presumed to be primitive relative to OCP1.

Here we reverse the evolution of the OCP by synthetically splitting the *Fremyella* OCP1 and OCP2 into their component domains and confirmed that the two polypeptides can interact and form heterocomplexes in the absence of the interdomain linker. The heterodimers derived from the domains of both OCP1 and OCP2 required carotenoid for heterodimer formation. Moreover, the heterodimers were photoactive. While the OCP1 rendered into its two component domains cannot re-associate *in vitro* after light-driven domain dissociation, the domains of heterodimeric OCP2 re-associate *in vitro* after photoactivation. The demonstrations of light-controlled heterodimer dissociation as well as the differences in re-association kinetics open up the possibility for engineering tunable photoswitches based on light-driven dissociation (and reassociation) of OCP domains. Moreover, we show that similar to NTD/RCP and HCPs (Wu and Krogmann, 1997; Kerfeld, 2004a; Leverenz et al., 2014; Leverenz et al., 2015; López-Igual et al., 2016), the CTD binds carotenoid on its own. Our data suggest that by forming a homodimer, two CTD polypeptides can effectively encompass the entire carotenoid. We refer to the CTD-Carotenoid homodimer as the C-terminal domain-like Carotenoid Protein (CCP). Collectively, these data support the domain fusion hypothesis for the evolution of the OCP and suggest that ancestral proteins for both the NTD and the CTD were likely to have been carotenoid-binding proteins. This implies that some members of the NTF2-fold protein superfamily can bind carotenoids, revealing a new family of soluble carotenoid binding proteins, with prospective members found across all kingdoms of life.

## Results

### Devolution of the OCP into a Heterodimer of the NTD and CTD

Given that the OCP is a structurally and functionally modular protein (Kerfeld et al., 2003; Leverenz et al., 2014) proposed to have evolved by the fusion of genes encoding homologs of its two constituent domains, we investigated whether the NTD and CTD could interact as separate polypeptides when synthetically isolated. Accordingly, the NTDs of the OCP1 and OCP2 from *Fremyella*, referred to hereafter as NTD1 and NTD2, respectively, were each fused to a C-terminal StrepTag, and the corresponding CTDs, CTD1 and CTD2, were fused to C-terminal HisTag (**Fig. 2A**). The cognate pairs of tagged NTD and CTD constructs were cloned together in different cloning sites of a single expression vector. These plasmids were expressed in *E. coli* strains with and without coexpression of genes for the biosynthesis of the ketocarotenoid canthaxanthin (CAN), and the resulting proteins were purified by affinity chromatography.

When the tagged NTD and CTD pairs were expressed in a CAN-producing strain of *E. coli*, orange-colored samples were obtained using tandem affinity purification followed by size-exclusion chromatography (SEC). These purified samples cross-reacted with either of the tag antibodies (anti-StrepTag or anti-HisTag) in Western blots (**Fig. 2B**), indicating successful pull-down of the two domains due to interaction. The cross-reacting bands migrated as 19.5 and 14.5 kDa in SDS-PAGE, which are the expected size of the individual tagged NTD and CTD domains, and CAN could be seen on blots of resolved heterodimers (orange bands at the bottom of **Fig. 2B**), suggesting release of the carotenoid upon denaturation with SDS. These results indicate heterodimer formation, in the presence of carotenoid, between the NTD and CTD of either OCP1 or OCP2.

In contrast, when the same plasmids were expressed in *E. coli* without CAN biosynthesis, colorless samples were purified with either HisTrap or StrepTrap affinity columns and showed elution of proteins corresponding to the size of the isolated domain constructs (**Fig. 2C**). These colorless samples only cross-reacted with antibodies for the corresponding affinity tag: no NTD was detected in peaks eluted from the HisTrap, and no CTD was detected in peaks eluted from the StrepTrap (**Fig. 2C**). Identical results were obtained when using either the OCP1 or the OCP2 from *Fremyella* as the source of the split and tagged domains. Thus, the isolated OCP domains were unable to interact in the absence of carotenoid, indicating that the carotenoid is required for heterodimer formation.

**Figure 2.**
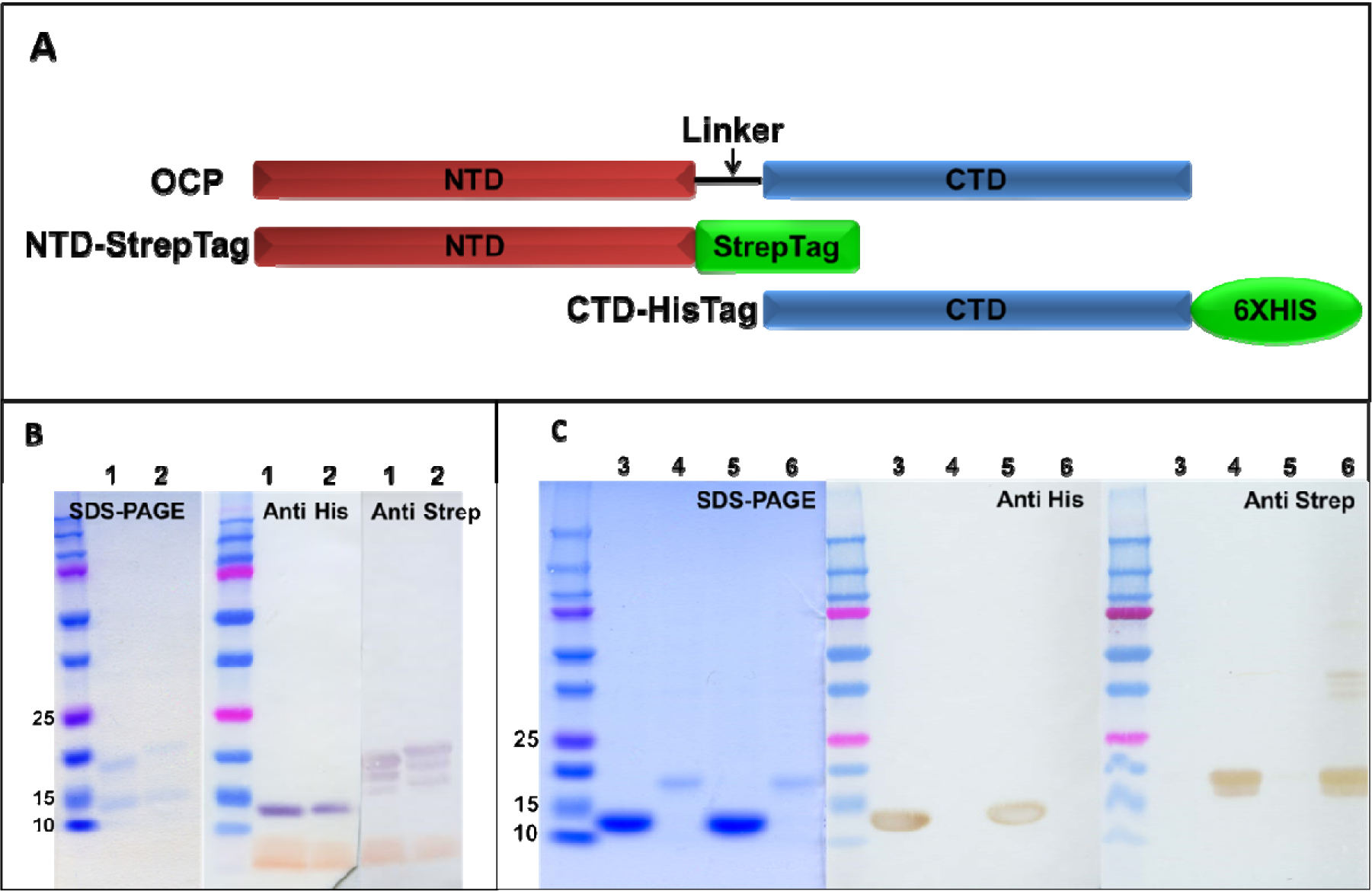
Cloning strategy and purification of NTD-CTD heterocomplexes from *E. coli.* **(A)** Domain architecture of the OCP, with experimental design shown for the tagged constructs used for expressing the isolated domains of OCP1 and OCP2 (NTD1, CTD1, NTD2, and CTD2) from *Fremyella* in this study. (B and C) SDS-PAGE and Western blot detected with anti-His and anti-Strep antibodies of purified NTD and CTD proteins from *E. coli*. **(B)** Proteins expressed in CAN-producing strain and purified by tandem affinity chromatography followed by SEC. Lane 1 - NTD1-CTD1; Lane 2 - NTD2-CTD2. **(C)** Same plasmids as in B were expressed in *E. coli*, without CAN production. Proteins were purified on Ni-column or separately on StrepTrap affinity column. Lane 3 –Results of the attempt to purify NTD1-CTD1 on Ni-column; Lane 4 - Results of the attempt to purify NTD1-CTD1 on StrepTrap column; Lane 5 – Results of the attempt to purify NTD2-CTD2 on Ni-column; Lane 6 – Results of the attempt to purify NTD2-CTD2 on StrepTrap column.

### Isolated CTDs Bind CAN

It is well established that the isolated NTD can bind carotenoid (Polívka et al., 2013; Leverenz et al., 2014; Leverenz et al., 2015); remarkably, we found that when only CTDs were expressed in CAN-producing *E. coli,* without the corresponding NTD, the purified proteins contained CAN (**Fig. 3A**). The absorption spectra of CTD1-CAN and CTD2-CAN showed a broad peak (140 and 141 nm peak-widths at half maximum for CTD1-CAN and CTD2-CAN, respectively) with no distinct vibronic features. These relatively broad peaks (when compared to the OCP^O^ spectrum) were similar in shape to the absorbance spectrum of the RCP (Wu and Krogmann, 1997; Kerfeld, 2004a; Leverenz et al., 2014; Leverenz et al., 2015), suggesting that the CAN molecule is not completely protected from solvent. The purified proteins were a mixture of apo-protein and holo-protein (compare A280 and A530 in **Fig 3D**). The yield of CTD1 expression was too low for further chromatographic analysis. While the CTD2 apo-protein appears in a monomeric (calculated size of 18 kDa) or dimeric form (calculated size of 37 kDa), the pigment is only observed in the CTD2 dimer (**Fig. 3D**), suggesting that the carotenoid is bound within a homodimer of two CTDs. We refer to the newly discovered CTD-CAN proteins as C-terminal domain-like Carotenoid Proteins (CCPs).

### Photoactivity of the Synthetic OCP Heterodimers

The absorbance maxima of the NTD1-CTD1 and NTD2-CTD2 heterodimers, purified in the dark by tandem affinity chromatography followed by SEC, are almost identical to the corresponding OCP-CAN maxima, with a slight difference in λmax for the shorter absorbance peak (2 nm blue-shifted peak for NTD1-CTD1 vs. OCP1, and 1 nm blue-shift for OCP2, **Fig. 3B**). Also, similar to the observations for the full-length OCP^O^s, the vibrational band resolution for NTD2-CTD2 is sharper relative to that of NTD1-CTD1 (**Fig. 3A** compared with Bao et al., submitted).

Both OCP1 and OCP2-based heterodimers are photoactive. Upon illumination, both photoactivate to the OCP^R^ form, as in full length OCP (**Fig. 3A**). In the red form, however, the λmax of both heterodimers are slightly red-shifted, compared to those of the full-length proteins (4 nm difference for OCP1, and 7 nm difference for OCP2, **Fig. 3B**). Additionally, in the red form the peak for the NTD2-CTD2 heterodimer was slightly broader (123 nm at half maximum) compared to the NTD1-CTD1 (119 nm), similar to the difference in peak width for full length OCP^R^s (Bao et al., submitted).

**Figure 3.**
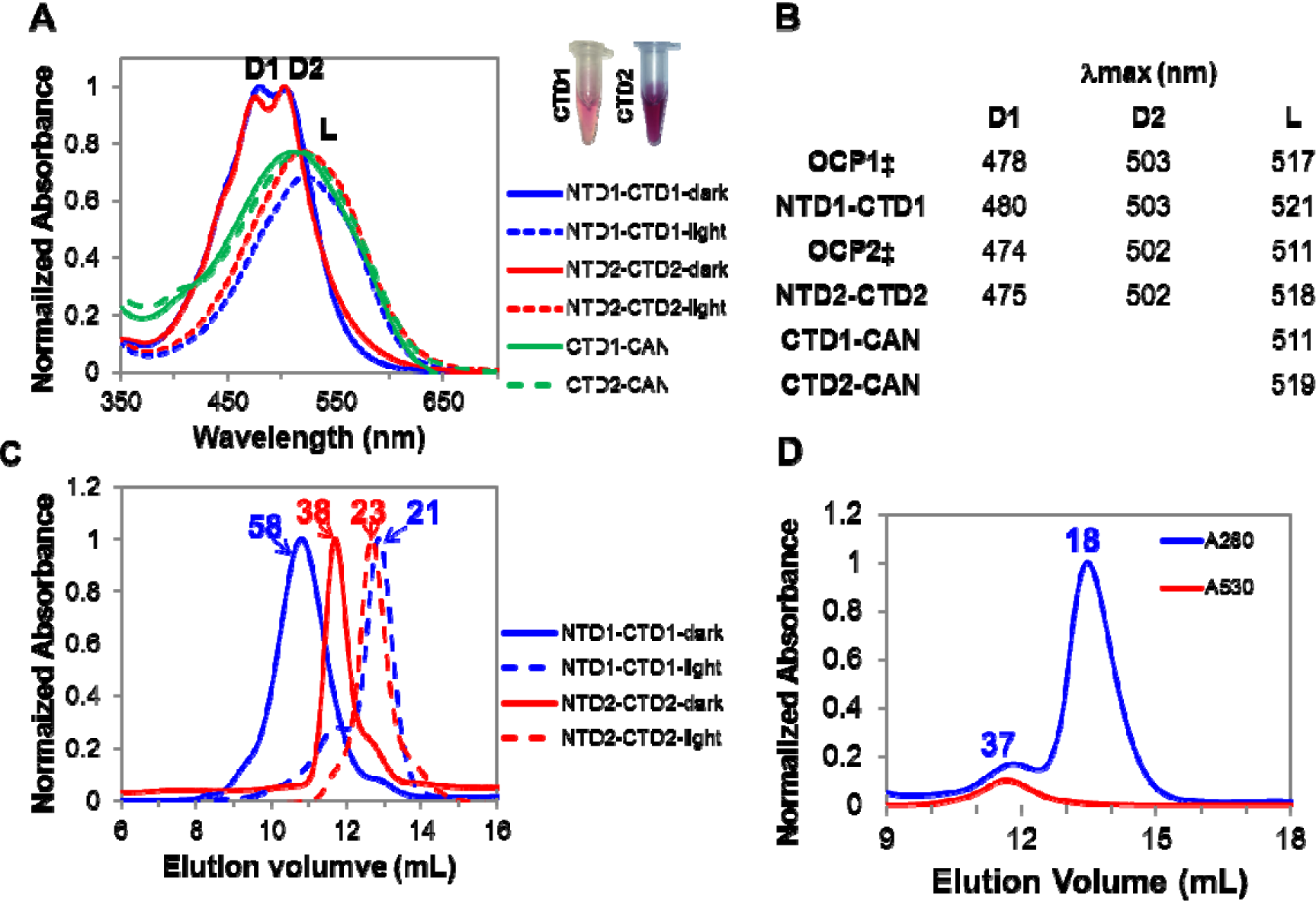
Biochemical characterization of split NTD1-CTD1 and NTD2-CTD2 heterocomplexes and CTD-CAN from *E. coli.* (**A**) Absorption spectra of heterodimers purified by SEC following tandem affinity chromatography on Ni column and Strep column, and CTD-CAN purified on a Ni column. Spectra of heterodimers purified in dark are represented by solid blue and red lines, and those of illuminated heterodimers are in dashed blue and red lines. NTD1-CTD1 (blue trace) and NTD2-CTD2 (red trace), CTD1-CAN (solid green) and CTD2-CAN (dashed green). Inset – Tubes with purified CTD1 and CTD2 expressed in the presence of CAN (**B**) Absorption maxima of purified NTD-CTD heterodimers and CTD1-CAN and CTD2-CAN from Panel A, compared to those of the corresponding full length proteins (**‡**) reported by Bao et al. (in preparation). “D1” and “D2” refer to the shorter and longer wavelengths of vibronic peaks observed in the dark OCP^O^–related form, respectively, and “L” refers to the absorption maximum of the photoactivated OCP^R^/RCP-related forms. (**C**) SEC of NTD-CTD heterodimers before and after illumination resolved on an analytical column. Trace colors same as in panel A. **(D)** SEC of CTD2-CAN purified on a Ni-column, protein absorbance at 280 nm (blue trace) and pigment absorbance at 530 nm (red trace) are shown. Calculated peak sizes in kDa are shown above the corresponding peaks on SEC.

We next examined the molecular masses of the heterodimers before and after illumination. In the dark, the NTD2-CTD2 heterodimer eluted in SEC at a volume corresponding to 38 kDa (**Fig. 3C**, solid red trace). As the expected size of the NTD2-CTD2 heterodimer is 35 kDa, comparable to monomeric, native OCP2, this suggests that the single heterodimer remains stable during chromatographic separation. In contrast, the NTD1-CTD1 heterodimer resolved at a calculated size of 58 kDa, significantly larger than the calculated 35 kDa size for the complex (**Fig. 3C**, solid blue trace), suggesting the presence of a dimer of heterodimers, comparable to the dimer formed by native OCP1s (Bao et al., submitted), from which these synthetic NTD1-CTD1 domains were derived.

Upon illumination of the NTD1-CTD1 and the NTD2-CTD2, the heterodimers dissociate into their component NTD and CTD (**Fig. 3C**, dotted lines). Once the heterodimers have been illuminated, both elute significantly later in SEC, performed at 4°C, with calculated sizes of 21 and 23 kDa for the dissociated NTD1-CTD1 and NTD2-CTD2 heterodimers, respectively. The calculated size of cloned NTDs is 19.5 kDa, and that of the CTDs is 15.5 kDa.

### Kinetics of Photoactivation and Domain Reassociation

Similar to full-length OCP1 and OCP2 from *Fremyella* (Bao et al., submitted), the kinetics of photoactivation and dark recovery of the heterodimeric OCPs are different. The NTD2-CTD2 heterodimer photoactivates more slowly upon illumination compared to the NTD1-CTD1 heterodimer (**Fig. 4A**, red vs blue traces, respectively), but recovery of the orange form was only observed with the NTD2-CTD2 sample (**Fig. 4B**, red traces). While the NTD1-CTD1 heterodimer converts from orange to red very quickly (T_1/2_ of 0.7 min at both 23 and 15°C), the NTD2-CTD2 heterodimer only photoconverts fully to the red form at lower temperatures (T_1/2_ of 2.9 min at 15°C). With elevated temperature, the NTD2-CTD2 photoconversion was incomplete (only 60%), albeit slightly faster (T_1/2_ of 1.7 min at 23°C). While the NTD1-CTD1 heterodimer did not recover in the dark under the experimental conditions (**Fig. 4B**, blue traces), the NTD2-CTD2 heterodimer recovered in the dark, with a faster rate at 23 °C than at 15 °C (T_1/2_ of 11.1 and 20.8 min at 23°C and 15°C, respectively).

**Figure 4.**
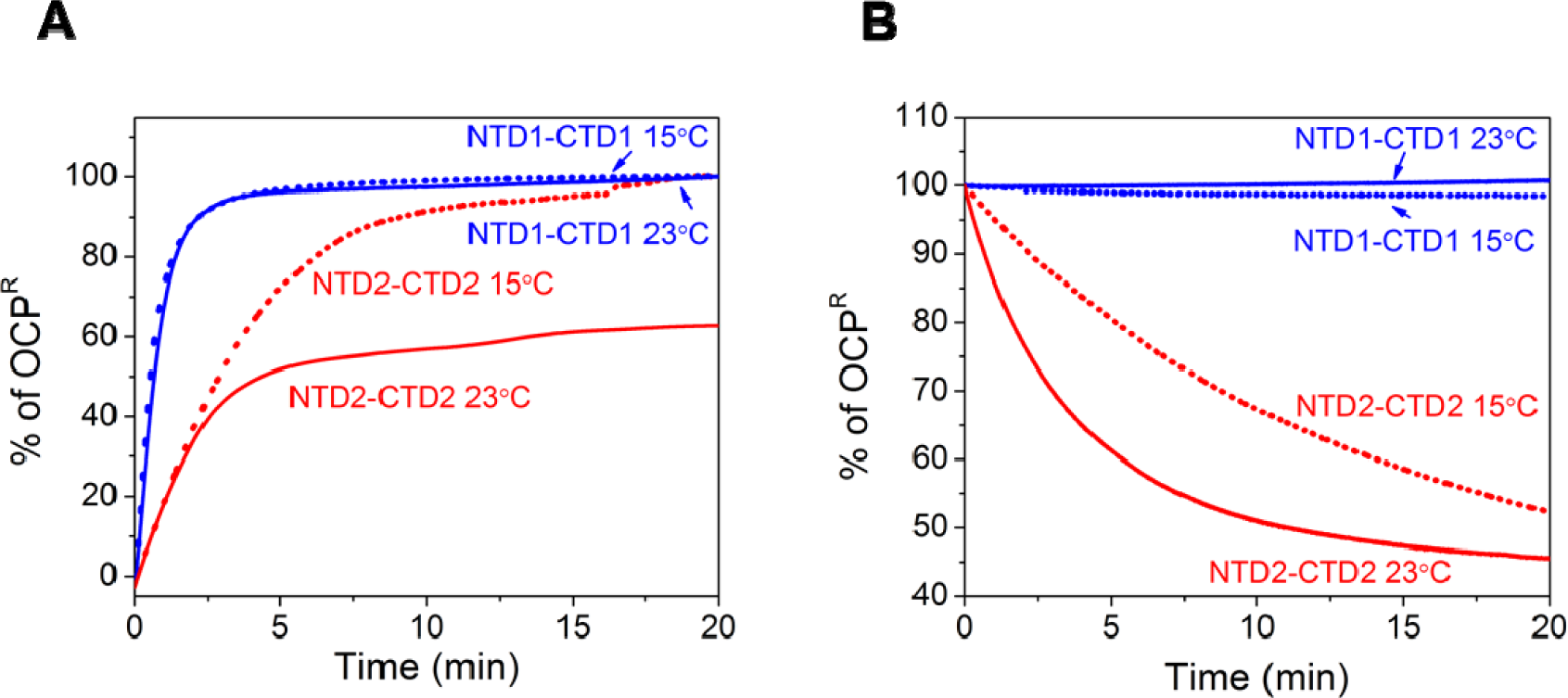
Kinetics of photoconversion and dark recovery of OCP heterodimers. (**A**) Kinetics of NTD-CTD orange to red conversion during blue-light illumination. (**B**) Kinetics of dark recovery of NTD-CTD heterodimers. Changes in the absorption were monitored at 550 nm at 23°C and 15°C for both experiments.

## Discussion

### Properties of OCP Heterodimers

Using the domains from either OCP1 or the OCP2 paralogs found in *Fremyella*, we demonstrate that covalent linkage is not required to form an OCP-like complex from its component domains, which interact when expressed as separate polypeptides in the presence of the carotenoid chromophore (Fig. 2). Strikingly, the NTD-CTD heterodimers produced from either family of the OCP are photoactive (Fig. 3, Fig. 4A), indicating that the interdomain linker is not required for photoactivation. Moreover, the requirement of CAN for stable NTD-CTD heterodimer formation in the dark (Fig. 2) supports the idea that the carotenoid itself is another module in the OCP system (Melnicki et al., 2016) and is involved in interdomain assembly of native OCPs. This firmly establishes the auxiliary and essential role of the pigment as a structural “bolt” which holds the two domains of the protein together in the dark (Cogdell and Gardiner, 2015) and that is capable of “un-bolting” the heterodimer upon photoactivation. Illumination causes domain dissociation and a 12 Å translocation of the pigment fully into the NTD (Gupta et al., 2015; Leverenz et al., 2015). The light-driven domain dissociation demonstrated here is in agreement with these findings (**Fig. 3C**). The finding that the carotenoid stays attached to the NTD (Leverenz et al., 2014), suggests that although the extant CTD has retained some carotenoid-binding capability, the affinity of the NTD to the carotenoid is stronger than that of the CTD.

Here we demonstrate for the first time that the CTD can bind carotenoid on its own as a homodimer (Fig. 3). This establishes unequivocally that the structural fold of the NTF2-like superfamily, to which the CTD belongs, is capable of serving as a carotenoid-binding domain. Moreover, this observation has provocative implications: some of the uncharacterized NTF2-like representatives in plants and other organisms may now be discovered to be previously-unrecognized water soluble carotenoid-binding proteins.

The absorbance spectra of the RCPs which result from NTD detachment upon photoactivation have a broad shape, lacking vibronic features; this is also observed for the CCPs resulting from CTD expression in the absence of NTD, suggesting that the carotenoid in the CCP is, as in the RCP, more solvent accessible than in the OCP. Although the structural details of carotenoid binding in the CCP remain uncertain, it is tempting to speculate that its CTD-CTD homodimer is symmetrical, with an elongated binding pocket spanning the two monomers, resembling how the OCP^O^ structure is bolted by the carotenoid spanning the NTD-CTD interface. In CCP, this could theoretically be formed by a second CTD monomer positioned where the NTD is located in OCP^O^. By interfacing to juxtapose the two carotenoid tunnels, the CAN could be largely shielded from solvent. Evidence supporting this is found in a dimeric crystal structure of a *Campylobacter* transpeptidase which is also a member of the NTF2 superfamily, [PDB:3K7C], where the ligand-binding cavities of two individual subunits combined to form a larger binding pocket (Eberhardt et al., 2013).

Full length OCP1^O^ from different organisms has been repeatedly shown to form dimers in crystals and in solution (Kerfeld et al., 2003; Wilson et al., 2010; Zhang et al., 2014; Liu et al., 2016; López-Igual et al., 2016; Bao et al., submitted). The finding that the NTD1-CTD1 heterocomplex elutes as a dimer from SEC (**Fig. 3B**) is consistent with these observations. In contrast, the NTD2-CTD2 heterodimer was only found as a monomer (**Fig. 3B**), in agreement with the findings for full length *Fremyella* OCP2 (Bao et al., submitted). The oligomeric state of the OCP has been hypothesized to play a regulatory role: photoactivation of OCP1 is proposed to first require dimer dissociation into monomers, as occurs with many enzymes (Marianayagam et al., 2004), before light-driven domain dissociation and carotenoid translocation can occur (Zhang et al., 2014; Bao et al., submitted). Dimeric interactions such as conserved salt-bridges observed in the crystal structures of full length OCP1^O^ (Kerfeld et al., 2003; Wilson et al., 2010; López-Igual et al., 2016) seem to persist in the synthetic NTD1-CTD1 heterocomplex, indicating that the linker is also not involved in OCP1 dimerization.

While dark recovery of the NTD2-CTD2 heterodimer was observed at both measured temperatures, the dark recovery of NTD1-CTD1 heterodimer was not observed at 23°C and was very slow at 15°C (**Fig 4B**). These results are in agreement with the recovery rates obtained for the corresponding full length proteins (Bao et al., submitted). It has been suggested that the linker might be required for dark recovery of *Synechocystis* OCP (Liu et al., 2016). There are no significant differences in sequence conservation between the linker sequences of OCP1s vs OCP2s (Bao et al., submitted). Moreover, the recovery of NTD2-CTD2 heterodimer in the absence of linker (**Fig. 4B**) proves that the linker is not fundamentally required for recovery. The inability of NTD1-CTD1 heterodimer to recover in the dark might be related to its requirement of FRP (Boulay et al., 2010); however, even when purified FRP was added to the reaction mixture *in vitro*, as shown for full-length OCP1 (Bao et al., submitted), the NTD1-CTD1 heterodimer did not recover under the experimental conditions.

The photoactivation kinetics of NTD1-CTD1 heterodimer is faster than that of NTD2-CTD2 (**Fig. 4A**). This is in contrast to the faster photoactivation of OCP2 compared to OCP1 (Bao et al., submitted). It is possible that the faster rate of orange to red conversion of NTD1-CTD1 heterodimer is a result of the inability of the heterodimer to recover under the experimental conditions (**Fig. 4B**). While the photoactivation rate of the NTD2-CTD2 reflects an equilibrium rate between photoactivation and dark recovery, the recovery of NTD1-CTD1 is negligible, resulting in an apparent faster rate of orange to red conversion. Additionally, the photoconversion of the NTD1-CTD1 heterodimer has weak to no temperature dependence, suggesting that under the experimental conditions, the kinetics depend only on light intensity and not temperature. This result is in contrast to kinetics of full-length OCP1, which show a significantly slower conversion rate from orange to red at 8°C, compared to 15°C (Bao et al., Submitted) and might, again, be related to the fact that there is no dark recovery in the synthetically split complex.

### The OCP Evolved from the Fusion of Two Primitive Carotenoid Binding Proteins

Gene recombination between protein domains often drives the evolution of new functions by fusion of previously interacting proteins into a single polypeptide, ensuring they are co-localized and co-expressed (Enright et al., 1999; Marcotte et al., 1999). Furthermore, multidomain proteins arising from a gene fusion event have been shown to frequently have gained a more specific or complex function compared to the homologous single-domain proteins from which they have been fused (Bashton and Chothia, 2007). In this manner, the OCP was hypothesized to have evolved as a result of a domain fusion event between genes encoding a carotenoid-binding HCP homologous to the NTD and a carotenoid-binding member of the NTF2-like superfamily homologous to the CTD (now referred to as CCP) (Kerfeld, 2004a; Melnicki et al., 2016). This domain fusion hypothesis is also supported by the frequent encoding of contemporary HCP4s and HCP5s at loci adjacent to a CTDH gene, (Kirilovsky and Kerfeld, 2013; Melnicki et al., 2016), a probable member of the CCP family.

We propose that such fusion between genes for an RCP-like HCP and an ancestral CCP-like protein (Fig. 5, step 3) would have altered the physiological implications of such an already-interacting pair in certain fundamental ways. For instance, forced co-localization of the CTD regulatory module and an HCP effector module could ensure its proximal availability, enhance turnover kinetics, constrain 1:1 stoichiometry between modules, prevent post-translational modifications (e.g., oligomerization) of the CCP, and/or enhance the affinity between domains (Enright et al., 1999; Marcotte et al., 1999). Duplication of this early ancestral form of the OCP would have enabled multiple subtypes to develop, distinguished by their different kinetic properties. Indeed some of these traits have been observed in this study, particularly for the OCP2-derived NTD2-CTD2 pair which appears to be more related phylogenetically to the ancestral “OCPx” subtype (Fig. 5, step 4) than the more advanced OCP1 subtype (Bao et al., submitted): kinetics are faster for full-length OCP2 compared to OCP1 (Bao et al., submitted), and NTD2-CTD2 interaction appears to be stronger (Fig. 4). It is plausible that the inability of CTD1 to reassociate with NTD1 after illumination and domain dissociation may be due to an evolutionary relaxation of domain interaction developed as a consequence of gaining the FRP mechanism to turn off quenching as a further step in the gain of regulation of OCP function (Fig. 5, step 5).

It is unknown whether photoactivation might have originated before or after the fusion of an HCP and a CCP into the primordial OCP (Fig. 5, step 2). The split OCP2 (but not OCP1) domains can reform the orange heterocomplex in the dark after photoactivation (**Fig. 4B**). Hence, based on shared affinity to the carotenoid ligand, the function of the CCP-like ancestor might have been to turn off quenching for an otherwise constitutively-active RCP/HCP4-like ancestor (Fig. 5, step 1) (Leverenz et al., 2014; López-Igual et al., 2016), similar to the dark recovery shown here for the NTD2-CTD2 complex. While the isolated, characterized HCPs are stably red, regardless of illumination (López-Igual et al., 2016), interaction and/or gene fusion with an NTF2-like domain (i.e. a CCP or a CTDH) likely enabled the evolution of both the photosensory and photoswitching ability. NTF2-like domains are found in some sigma factors involved in sensing environmental factors, and it has been suggested that such NTF2-like domains might be involved in light sensing in regulatory proteins (Aravind et al., 2010). Alternatively, as many NTF2-like containing enzymes such as ketosteroid isomerase, scytalone dehydratase, and phenazine biosynthesis protein B act on ketolated hydrophobic substrates, perhaps CCP interaction with HCPs arose via affinity for the ketocarotenoid, thus “pulling” it into the orange configuration and stabilizing the heterodimer.

**Figure 5.**
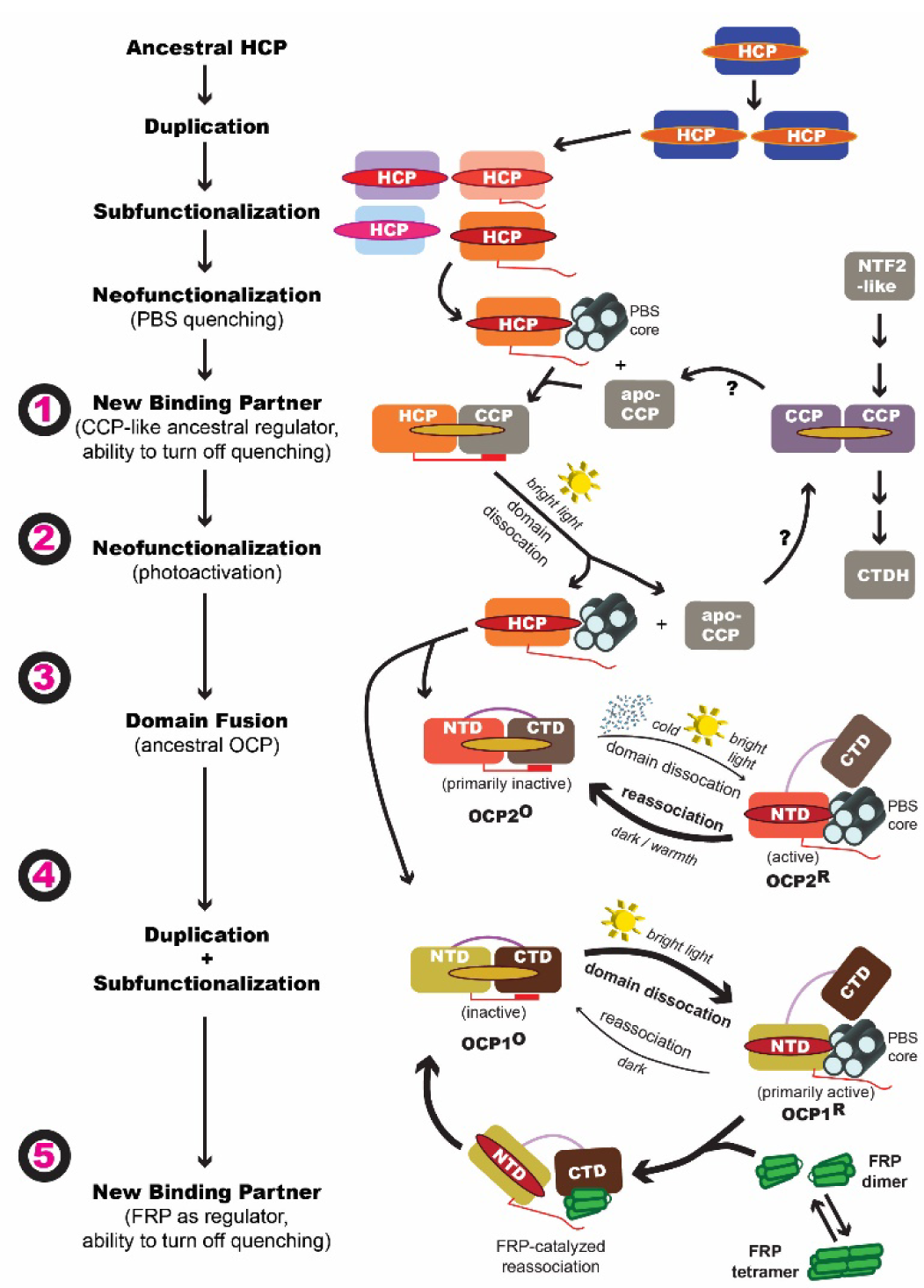
A model for OCP evolution. The evolution of the modern OCP is proposed to be the result of fusion of a particular HCP subtype resembling the RCP with an ancestral CCP-like homolog of the modern CTD capable of turning off quenching (1) by pulling the carotenoid between the two subunits, which could be reversed (2) by light-driven domain dissociation. Upon domain fusion (3), early OCPs would have prevented unnecessary quenching by maintaining a tight interaction between the fused domains (4), and photoactivating only under particular conditions such as bright light or cold. Later, this interaction became relaxed upon association with the FRP (5), broadening the dynamic range of quenching states available at any given time and relying on FRP and light intensity for fine-scale control of the quenching process.

### Implications for Biotechnological Applications

Domains are the structural, functional, and evolutionary units of proteins. Likewise, they can also be considered units of protein engineering (Kerfeld, 2015). Our findings provide the foundation to begin to use the modular structure of the OCP for optogenetic applications. The light-driven domain dissociation shown here (**Fig. 3C**), and the ability of the domains to reassociate (**Fig. 4B**), open the possibility for using the OCP for designing light-controlled applications. A first attempt in showing activity in a synthetic system was recently reported (Andreoni et al., 2017). Here, we have shown here that the heterodimeric OCP complexes retain photoactivity *in vitro*, even when affinity tags are fused to their C-termini. This suggests that it would be possible to fuse other proteins/cargo to the NTD and CTD domains without disrupting their ability to associate and disassociate.

Our results also underscore that the carotenoid is a third module in the assembly of native or devolved OCPs. It is not only involved in quenching PBS excitation and/or singlet oxygen, via the NTD, but also serves as a structural “bolt” to selectively hold together the domains in the dark. Altering the particular carotenoid – via expression in different carotenoid-producing strains, or by altering amino acids that comprise the binding pockets – has been demonstrated to affect the absorption properties (Melnicki et al., 2016), and thus may be exploited for “tuning” the sensor function to respond to different wavelengths and/or favor different targets for carotenoid delivery. Current designed photoswitches utilize other chromophores such as the flavins in LOV domains and bilins in phytochrome-based photoswitches (for example, Moglich and Moffat, 2010), but so far, no carotenoid-based photoswitch has been described. Adding carotenoid to the available chromophores used in synthetic biology would increase the spectral range of light that can be used in such applications (Mathes, 2016). Additionally, most currently engineered photoswitches that are designed to promote protein-protein interaction are based on light-driven domain association using phytochrome-based or LOV domain proteins (for example, Toettcher et al., 2011; Guntas et al., 2015), while an OCP-based engineered photoswitch would be based on light-driven domain dissociation, resulting in opposite effects.

The T_1/2_ time of dark recovery for the dissociated NTD2-CTD2 (11 minutes at room temperature) is significantly shorter than that of several LOV domain proteins such as the fungal protein vivid (VVD) (T_1/2_ 300 min), YtvA (50 min) and LOVK (~100 min) (Losi et al., 2003; Purcell et al., 2007; Zoltowski et al., 2009). With carefully targeted point mutations, the kinetics of some of these proteins have been modified and significantly accelerated (for example, Zoltowski et al., 2009), allowing more flexible targeting for optogentics designs. A similar approach can be used in order to engineer effective OCP-based photoswitches with faster kinetics or with biased rates favoring recovery over photoactivation. Our results suggest that perhaps an NTD1-CTD1-based platform can be used for applications requiring a “kill switch” for the function of the interacting target domains, as regulated by permanent domain dissociation following photoactivation; alternatively an NTD2-CTD2-based platform could be used for applications requiring more dynamic behavior consisting of either repetitive cycles of domain dissociation/reassociation or a dose-dependent response for “dialing” in the magnitude of the target function according to controllable LED light intensity or programmable flash cycles.

Application of an OCP-based photoswitch *in vivo* would have to be performed in an organism which produces ketocarotenoids naturally, or by heterologous expression of carotenoid biosynthetic genes in a similar manner to the *E. coli* strains used in this study (Bourcier de Carbon et al., 2015). Although only one group of plants is known to produce ketocarotenoids (Cunningham and Gantt, 2005), the process of converting nutritional yellow carotenoids to red ketocarotenoids (unexpectedly by a P450 homolog) has been recently established in birds (Lopes et al., 2016; Mundy et al., 2016), expanding the range of organisms in which utilizing an OCP-based photoswitch might be easier to implement. Furthermore, the revelation of carotenoid binding in the CTD, a member of the widespread NTF2-like protein superfamily, further extends the range of possible applications of photoswitches designed based on OCP architecture.

In summary, here we show for the first time that independently-expressed OCP domains can interact to form a stable heterocomplex capable of light-driven domain dissociation. These results support the hypothesis that the OCP has evolved through gene fusion of two different previously-interacting carotenoid-binding proteins. Our findings also demonstrate that the OCP domains are poised to become an optogenetic platform for a range of synthetic biology applications requiring controllable properties such as targeted spatial localization, tunable activity rates, or a functional “kill switch” using light. This engineering strategy of splitting the two OCP domains is essentially OCP devolution. We also demonstrate that the CTD represents a new family of carotenoid binding proteins within the large NTF2-like superfamily, which we name CCP. Given that members of the NTF2-like superfamily are found across all kingdoms of life, this revelation that the NTF2 fold can accommodate a carotenoid raises the possibility that the previously unrecognized CCP family may include novel carotenoprotein representatives in diverse organisms, beyond the cyanobacteria.

